# Evolutionary algorithms accelerate *de novo* design of potent Nectin-4-specific cancer biologics

**DOI:** 10.64898/2026.03.04.709551

**Authors:** Elisabeth Tan, Sophie C. Frechen, Bianca Broske, Julia M. Messmer, Tim N. Kempchen, Benjamin A. McEnroe, Dominic Ferber, Leonie Waluga, Eduard Below, Alexander Hoch, Michelle C. R. Yong, Johannes Oldenburg, Heiko Rühl, Matthias Geyer, Marieta I. Toma, Niklas Klümper, Tobias Bald, Jonathan L. Schmid-Burgk, Michael Hölzel, Gregor Hagelueken

## Abstract

Recent advances in AI-based structural biology have made *de novo* protein binder design increasingly effective, yet performance remains highly target dependent. For instance, the cancer surface antigen Nectin-4, an immunoglobulin-like cell adhesion protein, proved particularly challenging for RFdiffusion-based minibinder generation, yielding substantially fewer high-quality candidates than related targets. To address this bottleneck, we integrated an evolutionary genetic algorithm (GA) with AI-driven design. GA selection with tunable stringency was coupled with diversification via partial diffusion or direct sequence editing, enabling efficient exploration of sequence-structure space and rapid enrichment of promising candidates. This AI-GA pipeline quickly produced large and diverse minibinder panels with very good *in silico* quality metrics and is compatible with inputs from multiple design algorithms. Pooled, large-scale experimental screening identified highly stable Nectin-4 minibinders with single-digit nanomolar down to subnanomolar affinities. Lead binders were further engineered into Nectin-4-specific flow cytometry detection reagents and potent bispecific T cell engagers, demonstrating functional activity beyond binding. Together, these results show that evolutionary refinement can unlock challenging targets and accelerate *de novo* protein design for next-generation cancer biologics.

## Introduction

The advent of artificial intelligence (AI) has heralded a new era in structural biology, enabling the prediction of protein structures and the *de novo* design of proteins with unprecedented precision and speed^1–8^. The capability to engineer small, soluble proteins that selectively bind to predefined epitopes on target proteins has emerged as a transformative tool in biomedical research^4–7^. This level of control holds particular promise for the development of next-generation therapeutics, notably in oncology, where targeted binding to tumor-associated antigens can yield highly specific anti-cancer agents with reduced off-target effects. Moreover, for certain antibody–drug conjugates (ADCs), administration at or near the maximum tolerated dose has been associated with enhanced therapeutic efficacy^9^.

Despite recent progress, the efficiency of AI-driven binder design remains highly variable, exhibiting a strong dependence on the choice of target protein and selected epitope^10^. Recent studies have revealed that the physicochemical and geometric properties of the target’s molecular surface, such as curvature, electrostatic potential, and hydrophobic patches, play a critical role in dictating design success rates^4,10^. Understanding and optimizing these dependencies is therefore crucial for accelerating the pipeline from *in silico* design to experimental validation.

Genetic algorithms (GAs), with their foundation in principles of biological evolution, offer a robust framework for tackling complex optimization challenges characterized by high-dimensional parameter spaces^11,12^. By iteratively selecting, recombining, and mutating candidate solutions, GAs can efficiently explore diverse regions of sequence and structural space, making them well suited to refine AI-generated binder candidates. In this study, we integrate GA-based optimization with state-of-the-art AI-based techniques to enhance protein binder design. We apply our methodology to Nectin-4, a cell-adhesion molecule overexpressed in different solid cancers and the target of the approved ADC enfortumab vedotin (EV)^13–16^. AI-based binder generation with the RFdiffusion pipeline proved particularly inefficient for Nectin-4, in contrast to structurally related targets such as PD-L1^4,10^. Here, we show that the combined AI-GA approach markedly accelerates *in silico* design of Nectin-4 minibinders. GA-based refinement of seed designs produced specific Nectin-4 minibinders with single-digit nanomolar and even subnanomolar affinities. Proof-of-concept functionalization as flow cytometry detection reagents and bispecific T cell engagers (TCE) further demonstrates the applicability of the approach. Together, these findings establish a versatile strategy for rational, accelerated therapeutic *de novo* protein engineering and development of new targeted cancer treatments.

## Results

### Nectin-4, a difficult target for the RFdiffusion algorithm

Human Nectin-4 (UniProt ID: Q96NY8) is a cell-adhesion molecule overexpressed on the surface of cancer cells, most prominently in urothelial carcinoma, and also in subsets of lung, breast, and pancreatic cancers^13,14,17^. Using the recently published protein design algorithm RFdiffusion^4^, we aimed to create minibinder proteins (AI-minibinder) that recognize Nectin-4. It is known that the success of AI-minibinder design correlates with certain surface properties of the protein, for instance the presence of hydrophobic patches as hotspots for binding (**Figure 1a**)^4,10^. The extracellular part of Nectin-4 consists of three Ig-like domains and we chose the Ig-like V-type domain (residues 32-148) as our design target for RFdiffusion. However, we noticed that, compared with our previous efforts to design AI-minibinders against the Ig-like V-type domains of PD-L1, CD276 and VTCN1^10^, the yield of designs with promising predicted aligned error (pAE) interaction scores was very low, despite the fact that these proteins are structurally similar to Nectin-4 (**Figure 1b**). pAE interaction scores inversely correlate with the experimental success of AI-minibinders, with especially high success rates for values below 5, based on our experience and that of others^4,10^. The Nectin-4 minibinder design runs yielded ∼50-fold fewer promising candidates than the PD-L1 runs, effectively requiring at least ∼50-fold more computation time to generate a comparable number of AI-minibinder designs for subsequent experimental testing. When designing Nectin-4 AI-minibinders without predefined hotspots, the success rate was even lower (pAE<5: 0% and pAE<10: 0.6%). This reduced efficacy of designing promising Nectin-4 minibinders likely reflects the fact that PD-L1, unlike Nectin-4, exposes a prominent hydrophobic surface patch that provides a well-defined hotspot for interface selection and binding (**Figure 1a**). Overall, these results highlight that the efficiency of RFdiffusion-based protein binder generation can vary widely even across structurally related target proteins.

**Figure 1:**
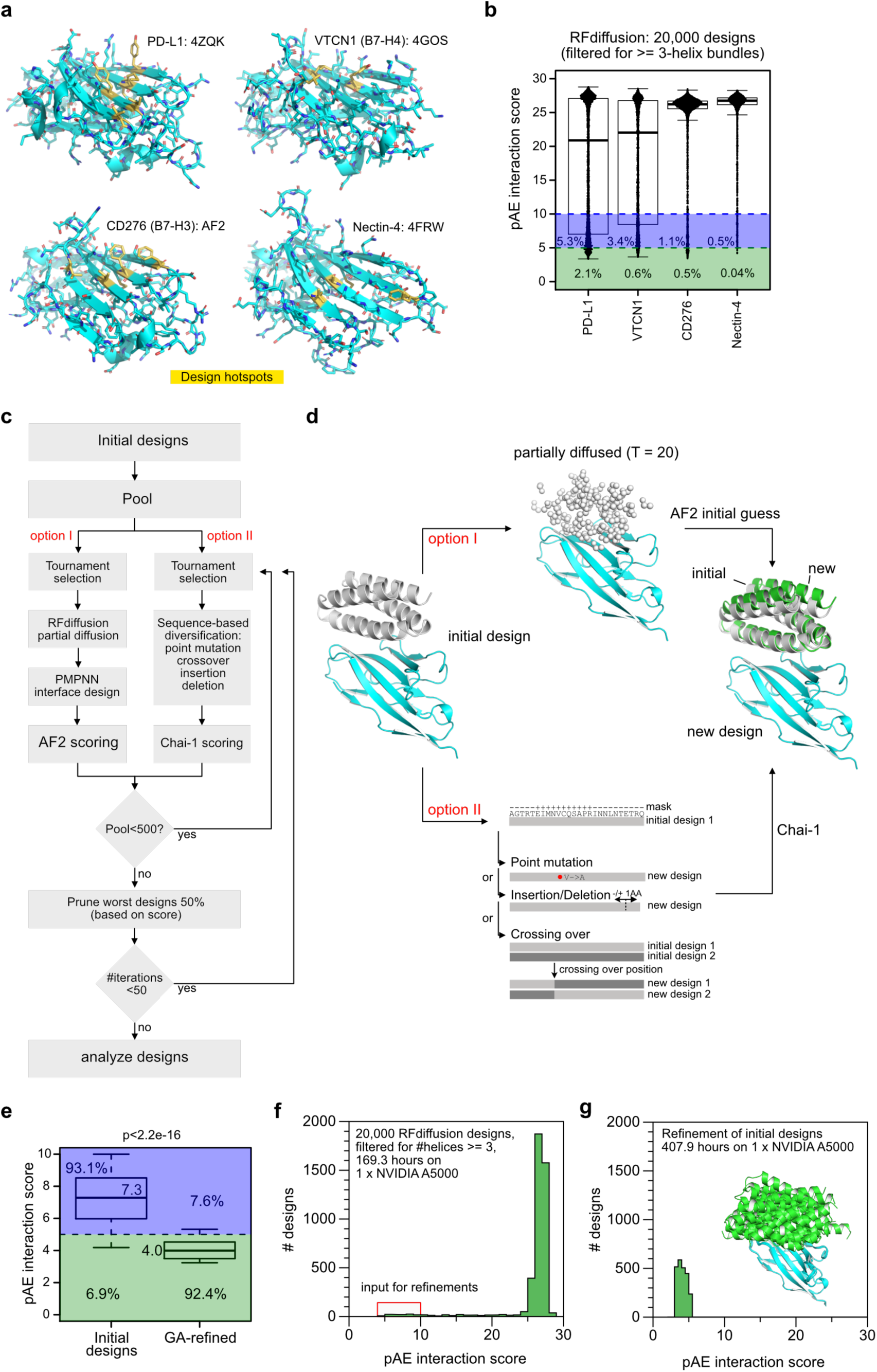
Efficient *in silico* refinement of Nectin-4 minibinders by genetic algorithm. **a)** Ig-like V-type domains of cancer cell surface proteins: PD-L1 crystal structure (PDB:4zqk; hotspots: I54, Y56, V68, M115, Y123), VTCN1 (B7-H4) crystal structure (PDB:4gos; hotspots: L72, L79, Y131), CD276 (B7-H3) AF2 model (hotspots: I66, L75, F123, F129), and Nectin-4 crystal structure (PDB: 4frw; hotspots: A66, L81, F132). **b)** Histograms summarizing the result of multiple RFdiffusion design runs for different targets. **c)** Block diagram presenting the evolutionary refinement algorithms described in this work. **d)** Schematic explaining the introduction of variations into initial AI-minibinder designs by either partial diffusion (option I) or sequence manipulation (option II). **e)** Boxplot showing pAE interaction scores of initial RFdiffusion designs and GA-refined (partial diffusion) designs. Median pAE scores are indicated and percentages of pAE<= 5 or pAE>5. Wilcoxon test. **f)** Histogram showing pAE interaction scores of the initial RFdiffusion design runs for Nectin-4. The designs that were used as inputs for the refinement algorithms are highlighted by a red box. **g)** Histogram showing the refinement results using option I of the evolutionary algorithm. Insert: Cartoon representation of Nectin-4 (cyan) and a set of diverse AI-minibinder designs (green).

### *In silico* refinement of AI-minibinder candidates by genetic algorithms

We wondered, whether the small number of promising designs could be refined or optimized by evolutionary algorithms to accelerate the process and increase the chances of success. Such algorithms are often used for difficult optimization problems with a large number of parameters and have been successfully applied to problems ranging from natural science to engineering^11,12^. In a nutshell, the process starts with the creation of trial solutions (here: AI-minibinder designs originating from RFdiffusion), which are scored to determine their “fitness” (for instance the pAE interaction or ipTM scores). The least fit members of this “population” are then pruned and the surviving designs are used to create offspring (i.e. new AI-minibinder designs), which are variations of their “parent” designs. A “tournament selection” is used to determine which designs are selected to create offspring: A number of designs are randomly picked and the fittest representative is used to create offspring. This procedure is repeated for multiple generations (cycles), ideally leading to an iterative optimization of the designs (**Figure 1c**). Such an evolutionary algorithm is also known as genetic algorithm (GA) or genetic cycle.

First, to validate the GA approach *in silico*, we used the outputs of multiple RFdiffusion runs with predefined interaction hotspots on Nectin-4 as input designs (**Figure 1a**). Benchmarked on a single NVIDIA RTX A5000 GPU, this required 169.3 hours of computation time. From a total of 20,000 AI-minibinder designs only 104 of these initial designs had a pAE interaction score < 10 (a common cutoff for promising candidates), and only 8 had a pAE interaction score < 5. In addition, we used 12 initial designs with a pAE interaction score < 10 (none < 5) out of 20,000 designs that we obtained when designing Nectin-4 AI-minibinders without predefined hotspots.

The variation of the initial designs can be achieved in different ways and we tested two options (option I and II) in this work (**Figure 1d**). Option I is based on the partial diffusion feature of RFdiffusion^4^. This feature can be used to add structural perturbations to the scaffold of the design, followed by the design of a new amino acid sequence using the ProteinMPNN algorithm (PMPNN)^18^. We applied this algorithm to the 104 and 12 best initial designs (seed designs) described above, and let the *in silico* evolution proceed for 50 generations. The procedure was then repeated for emerging top candidates in three independent runs, since genetic algorithms are prone to be trapped in local minima and repeated runs are a way to alleviate this problem. After a total calculation time of 407.9 hours on a single A5000 GPU, the algorithm had created hundreds of refined designs with very promising pAE interaction scores almost all (92.4%) below 5 (**Figure 1e-g**). Using the yield of designs with an interaction pAE<5 as the benchmark, GA-based refinement was ∼90-fold more compute-efficient than the conventional approach for obtaining the same number of promising designs. Due to our design criteria (size<= 88 amino acids and helical), all designs had a helical bundle fold (>= 3 helices) but were nevertheless structurally diverse (**Figure 1g**). The size restriction to 88 amino acids is driven by technical and economic constraints, as the coding sequence together with flanking primer regions must fit within 300-bp DNA oligo pools for cloning, and the helical-bundle criterion was imposed to enhance minibinder stability^10^.

We reasoned that AI-minibinder refinement could potentially also work in a manner more similar to a classical genetic algorithm by producing design variants not by partial diffusion, but simply by introducing point mutations, insertions, deletions or crossing-over operations to the amino acid sequences of a set of AI-minibinder candidates (**Figure 1d**). Here, the variability can be easily constrained to regions of interest and it is easy to control the types of amino acids that are introduced, defining thereby also the biophysical parameters of the designs. To implement this feature, we made use of the Chai-1 algorithm with embedding of the ESM2 protein large-language model^6^. The second option also led to AI-minibinder designs with very good scores as shown in the **Supplementary Figure 1**. It should be noted that the fitness parameter was in this case based on an adjustable linear combination of the pTM score of the minibinders and the ipTM score of the minibinder-target complex to avoid designs that fit perfectly to the target epitope but at the cost of an instable fold (**Supplementary Figure 1b-d)**. Furthermore, imposing additional constraints in this algorithm, such as a desired target isoelectric point (pI), hydrophobicity (GRAVY score) or epitope, is possible, but comes at the expense of efficient optimization of the ipTM score (**Supplementary Figure 1e**).

### Experimental validation of refined AI-minibinder designs

The following experiments focus on the experimental evaluation of the seed designs (n = 116) and their refined offspring designs (n = 1876) that were created with option I (**Figure 1c**). To identify functional AI-minibinders, we used our previously described mammalian cell-surface display pipeline^10^. Briefly, we encoded each AI-minibinder in a chimeric antigen receptor-like construct, allowing its presentation on the surface of HEK293T cells (**Figure 2a**). Cells bearing functional AI-minibinders were detected by staining with human Nectin-4 Fc fusion proteins and isolated via fluorescence-activated cell sorting (FACS). The DNA of the enriched population was extracted, and the corresponding AI-minbinders were identified by deep sequencing showing a success rate of 4.2% (**Figure 2b**, left panel). Importantly, all functional AI-minibinders emerged during genetic algorithm (GA) refinement, whereas none of the original seed designs were recovered.

**Figure 2:**
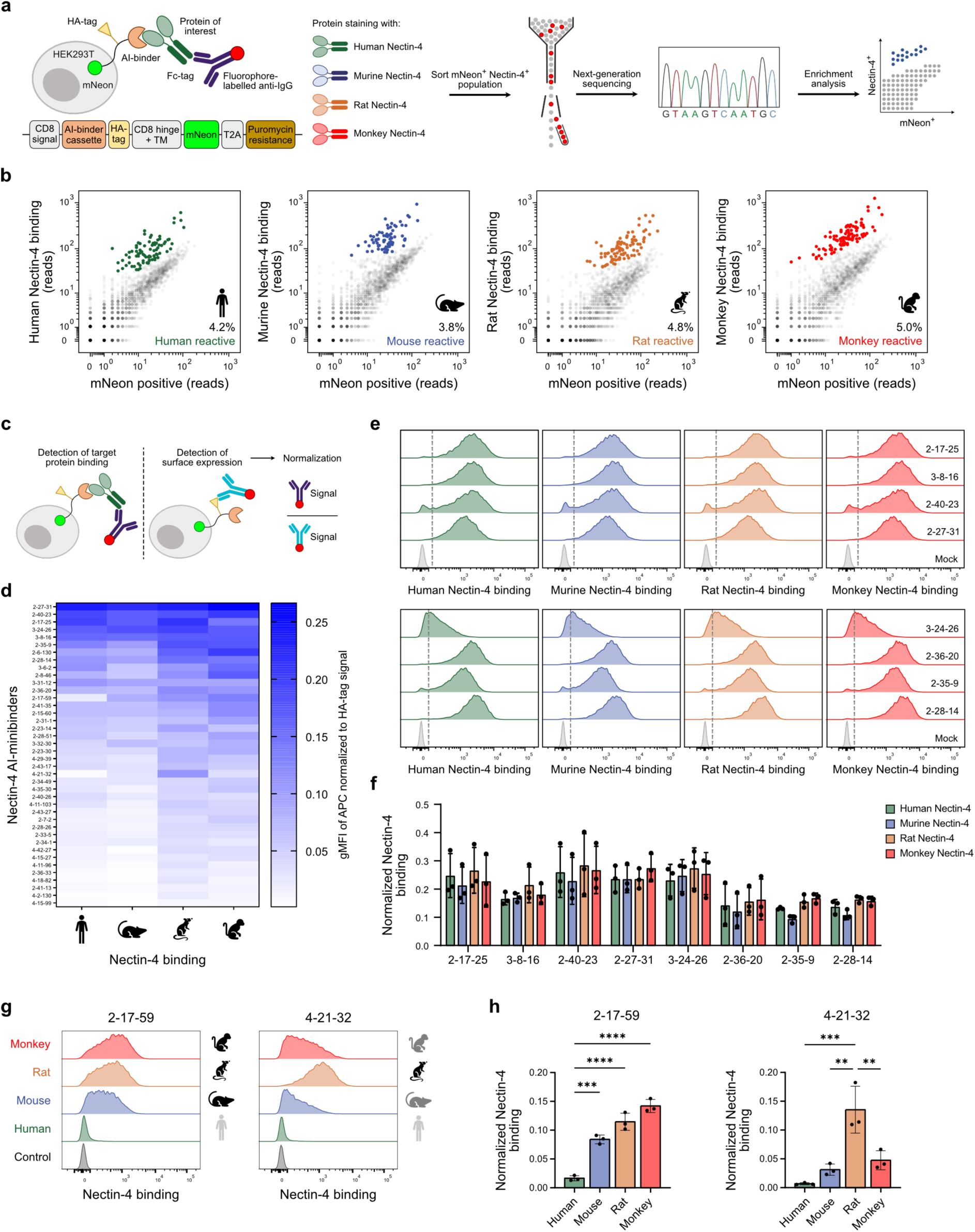
Experimental identification of functional Nectin-4 AI-minibinder designs. **a)** Scheme illustrating the experimental screening pipeline using the mammalian cell-surface display for AI-minibinder presentation, fluorescence-activated cell sorting and next-generation sequencing to identify candidates. **b)** Scatter plots showing enriched minibinders for human, murine, rat and monkey Nectin-4 (colored dots). **c)** Cartoon showing the experimental set-up of surface presentation of the AI-minibinder on HEK293T cells and detecting the binding to the target protein. Staining of the HA-tag was used to normalize the protein binding to the surface expression of the AI-minibinder. **d)** Heatmap summarizing the binding to human, murine, rat and monkey Nectin-4 Fc-fusion proteins of 40 individually validated Nectin-4 AI-minibinders using HEK293T cell-surface display. Data are shown as geometric mean fluorescence intensity (gMFI) normalized to the surface expression detected with the HA-tag signal. **e)** Representative histograms showing flow cytometry analyses of the binding to Nectin-4 Fc-fusion proteins from the different species for 8 AI-minibinder candidates. **f)** Quantification of the experiment described in e. **g)** Representative histograms of minibinder 2-17-59 showing selective binding to Nectin-4 Fc-fusion protein from murine, rat and monkey origin (left) and minibinder 4-21-32 demonstrating preferential binding to rat Nectin-4 Fc-fusion protein (right). **h)** Quantification of the experiment described in g for binder 2-17-59 (left) and 4-21-32 (right). Data are shown as mean ± SD performed in biological triplicates (n = 3). **p<0.01; ***p<0.001; ****p<0.0001; two-way ANOVA with Tukey’s multiple comparisons test.

To investigate the cross-species reactivity, we repeated the screen with homologs of Nectin-4 from mouse (92% identical amino acids to human Nectin-4), rat (83.9%) and monkey (99.4%). Of note, success rates were comparable for murine (3.8%), rat (4.8%) and monkey (5%) Nectin-4 (**Figure 2b**). As shown, most of the identified AI-minibinders seemed to bind Nectin-4 from all four species, which may not be surprising, due to the high evolutionary conservation of the target protein. However, some candidates that bound to the murine protein did not bind the human protein and *vice versa*. The same observation was also made for the other orthologs (**Supplementary Figure 2**). Next, we selected 40 AI-minibinders and validated Nectin-4 binding in individual flow cytometry assays, with signals normalized to AI-minibinder surface expression (**Figure 2c**). Consistent with the pooled screening results, we experimentally validated that the majority of the AI-minibinders showed binding to Nectin-4 across all four species (**Figure 2d-f**). Only a few AI-minibinders revealed deviating binding patterns with poor interaction to human Nectin-4 (**Figure 2g, h**).

### Biochemical and biophysical characterization of AI-minibinders targeting human Nectin-4

Based on the data above, the eight best AI-minibinder candidates were selected for a thorough biophysical characterization. The AI-minibinders were cloned into the pEHISTEV vector^19^ and expressed in *Escherichia coli*. All candidates showed high expression levels and were purified to homogeneity via His-tag affinity purification and size exclusion chromatography (SEC) (**Figure 3a, b**).

**Figure 3:**
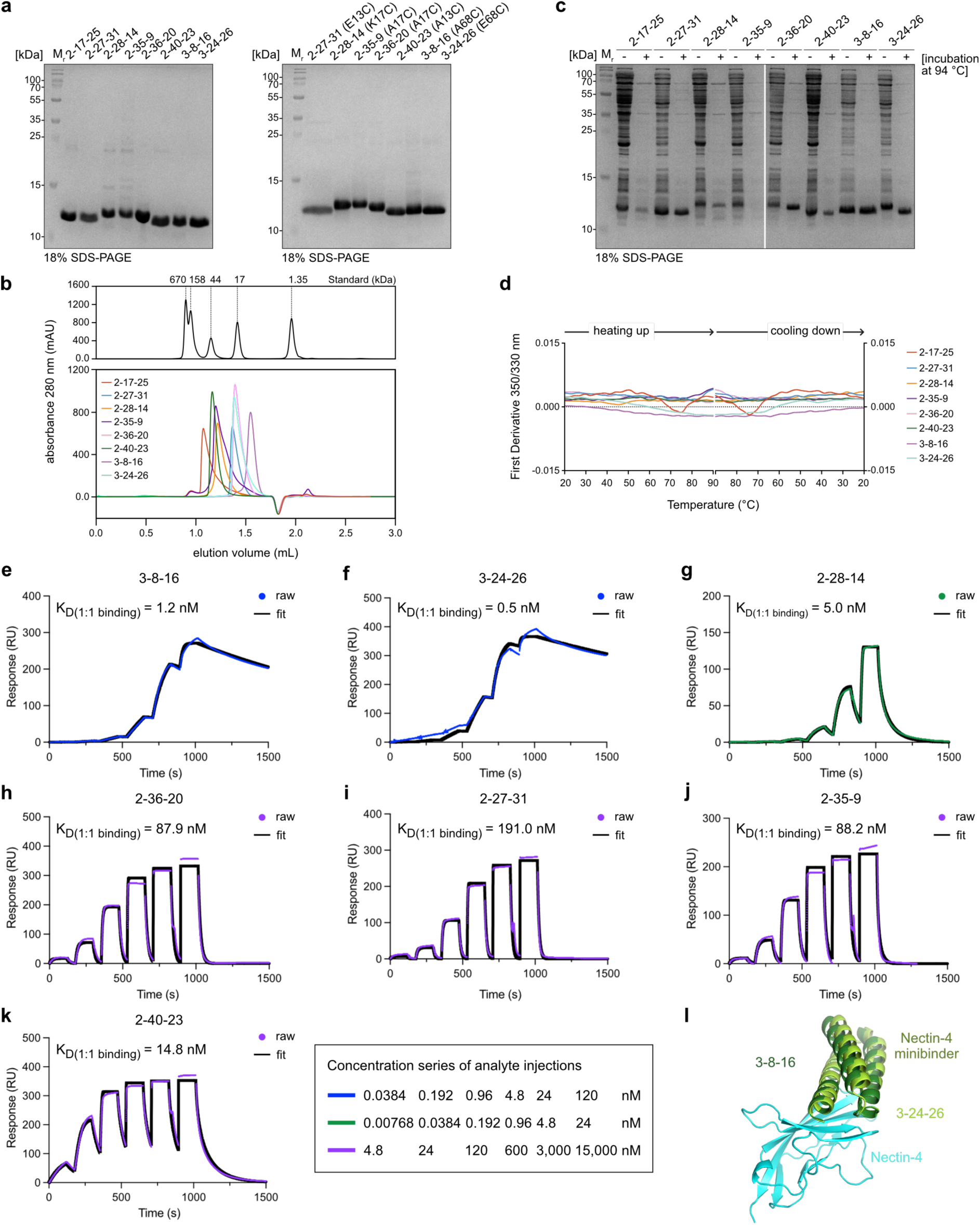
Biochemical characterization of Nectin-4 AI-minibinders. **a)** Coomassie-stained SDS-PAGE of purified Nectin-4 AI-minibinder proteins that were overexpressed in *E. coli*. Left: original designs, right: biotinylated versions of minibinders including an engineered mutant cysteine residue on the ‘backside’ of the AI-minibinder, opposite of the interaction interface. **b)** Analytical SEC of purified AI-minibinder proteins using a Superdex Increase 75 3.2/300 column. SEC traces of a commercial standard are included as molecular weight reference. **c)** Coomassie-stained SDS-PAGE of *E. coli* supernatants after the overexpression of AI-minibinders with (+) and without (−) incubation at 94 °C for 20 min. The boiled samples were centrifuged and the cleared supernatant was loaded onto the gel. **d)** Nano differential scanning fluorimetry (nanoDSF) analysis showing the thermal stability of the AI-minibinder proteins. The y-axis shows the first derivative of the corresponding melting curve. The two panels show the unfolding (left) and refolding (right) phases of the experiment. **e-k)** SPR (surface plasmon resonance) sensorgrams showing raw binding data (colored dots) and fitted curves (black) used to determine binding affinities (K_D_) of Nectin-4 AI-minibinders. The concentration series of analyte injections are indicated in the legend. **l)** Structural model of 3-8-16 and 3-24-26 Nectin-4 minibinders (shades of green) in complex with Nectin-4 (cyan) as determined in Chai-1 shown as overlay.

Interestingly, the elution profiles suggested monomeric (3-8-16), and oligomeric forms (dimer to tetramer) of the AI-minibinders but did not show signs of uncontrolled aggregation. Some SEC traces showed trailing peaks, indicating that the observed oligomerization is concentration dependent. As shown in **Figure 3c-d**, the proteins had exquisite thermal stability, a feature that has been previously described for AI-minibinders^10^. We used surface plasmon resonance (SPR) spectroscopy to determine the binding kinetics and affinities of the AI-minibinder proteins towards human Nectin-4. We immobilized the target protein to the sensor surface and injected a concentration series of the recombinant AI-minibinders. The sensorgrams shown in **Figure 3e-k** clearly show strong interactions between the AI-minibinders and the target protein. Many of the determined K_D_ values were in the low nanomolar range, with the highest affinities of approximately 1.2 nM and 0.5 nM observed for AI-minibinders 3-8-16 and 3-24-26, respectively (**Figure 3e, f**). Both minibinders are closely structurally related to each other (**Figure 3l**). Notably, all interactions were fitted best under the assumption of a 1:1 binding mode, conforming mono-dispersity of the AI-minbinders. As reported in our previous study, the binding kinetics varied strongly among the different designs, ranging from very fast to very slow on- and off-rates. This confirms that it is beneficial to test a larger number of AI-minibinder designs and select those designs that fit best to the target application.

### Specificity of Nectin-4 quattrobinders in flow cytometry

We had previously shown that AI-minibinders can be rapidly converted into antibody-like detection reagents for applications such as flow cytometry^10^. Here, we engineered a single cysteine residue into the non-binding surface (’backside’) of our AI-minibinders and assembled them with AF647-labelled streptavidin to generate tetravalent detection reagents termed quattrobinders (**Figure 4a**). Note that the cysteine version of 2-17-25 did not express well and was therefore not included in the following experiments. **Figure 4** illustrates the performance of the Nectin-4 quattrobinders in flow cytometry across multiple cell lines.

**Figure 4:**
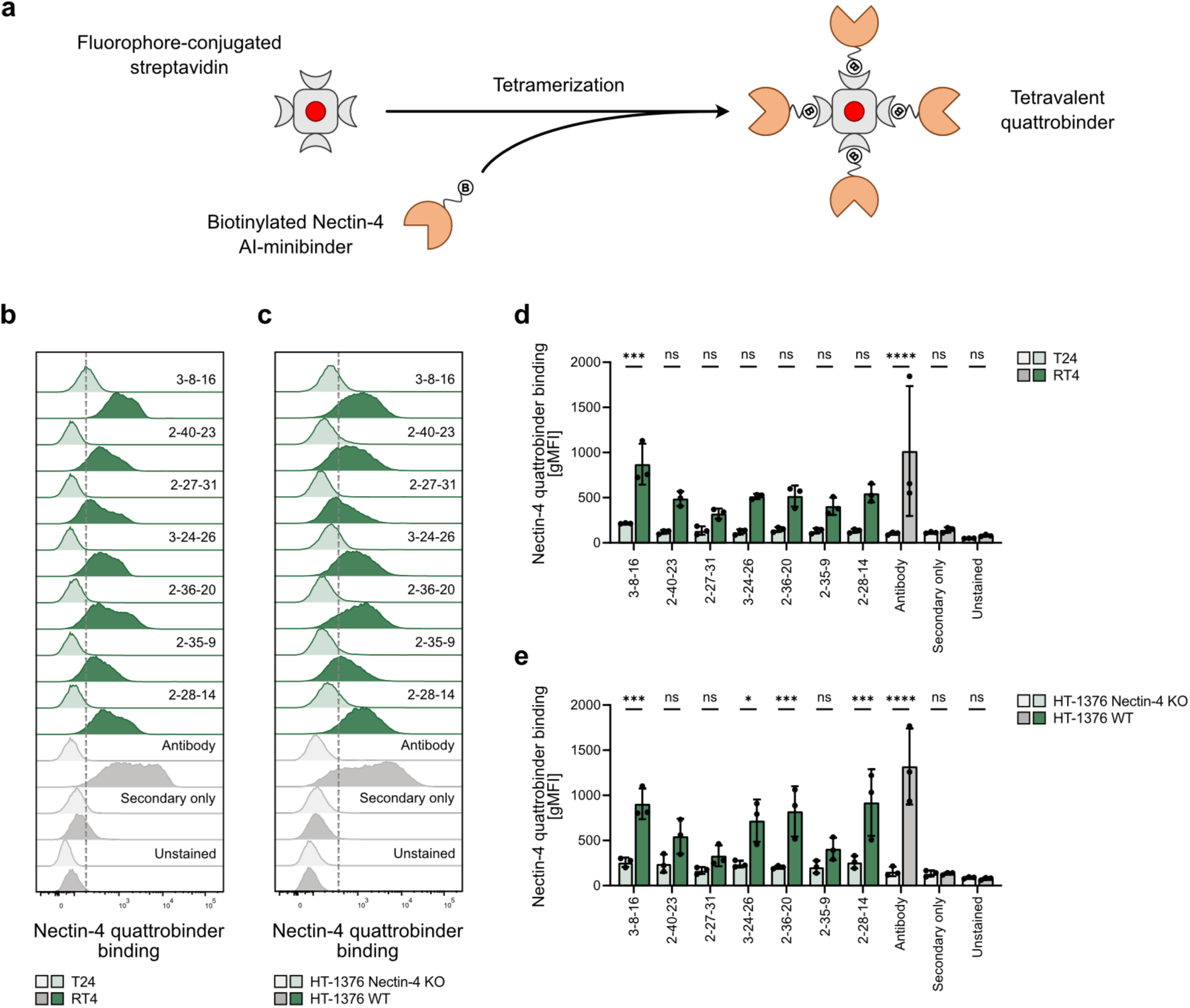
Nectin-4 AI-minibinders detect endogenously expressed Nectin-4 on cancer cells as multivalent quattrobinders. **a)** Schematic illustration of the assembly of biotinylated Nectin-4 AI-minibinders with fluorophore-conjugated streptavidin as tetravalent quattrobinder. **b-c)** Representative histograms showing flow cytometry analyses of AF647-conjugated Nectin-4 quattrobinders in comparison to a conventional biotinylated Nectin-4 antibody with a secondary AF647-streptavidin staining on target expressing urothelial cancer cells (RT4) and non-expressing cancer cells (T24) as well as on HT-1376 WT and the corresponding knockout cell line (HT-1376 Nectin-4 KO). **d-e)** Quantification of the experiment described in b and c. Data are shown as mean ± SD performed in biological triplicates (n = 3). ns – non-significant; *p<0.05; ***p<0.001; ****p<0.0001; two-way ANOVA with Šídák’s multiple comparisons test.

Nectin-4 is highly expressed on luminal urothelial carcinoma (UC) cells such as RT4, but is low or absent on basal-type UC cells such as T24. To benchmark our quattrobinders, we included a well-characterized Nectin-4-specific monoclonal antibody commonly used for flow cytometry^14^. All Nectin-4 quattrobinders robustly stained RT4 cells, whereas signals on T24 cells remained at background levels, closely mirroring the staining pattern of the reference antibody (**Figure 4b, d**). Similar results were obtained when staining Chinese hamster ovarian (CHO) cells overexpressing human Nectin-4 compared to parental CHO cells (**Supplementary Figure 3a, b**). Among the constructs, quattrobinder 3-8-16 performed best, consistent with its favorable biophysical profile and high affinity (K_D_ = 1.2 nM). To further assess specificity, we used a polyclonal *NECTIN4* knockout (KO) culture of HT-1376 cells using CRISPR-Cas9 (**Supplementary Figure 3c, d**)^14^. HT-1376 is a luminal type UC cell line that harbors amplifications of the *NECTIN4* gene^13^. Again, the Nectin-4 quattrobinders stained HT-1376 wild-type (WT) cells, while signals on HT-1376 Nectin-4 KO cells were reduced to background, as independently confirmed with the reference antibody (**Figure 4c, e** and **Supplementary Figure 3e**). Collectively, these data show that Nectin-4 AI-minibinders not only bind recombinant Nectin-4 *in vitro* with very high affinity, but also detect endogenous Nectin-4 specifically on the surface of UC cell lines, when formatted as quattrobinders for flow cytometry.

### Constructing Nectin-4 bispecific T cell engagers

Next, we aimed to functionalize the Nectin-4 AI-minibinders into biologics, and as proof-of-concept, we aimed to create bispecific T cell engagers (TCEs) (**Figure 5a**). In such constructs^20,21^, the well-described OKT3 single-chain variable fragment (scFv) binds CD3ε on T cells^22,23^ mediating T cell engagement. The second binding arm recognizes a highly expressed cancer cell surface antigen and is therefore responsible for target cell engagement (**Figure 5a**).

**Figure 5:**
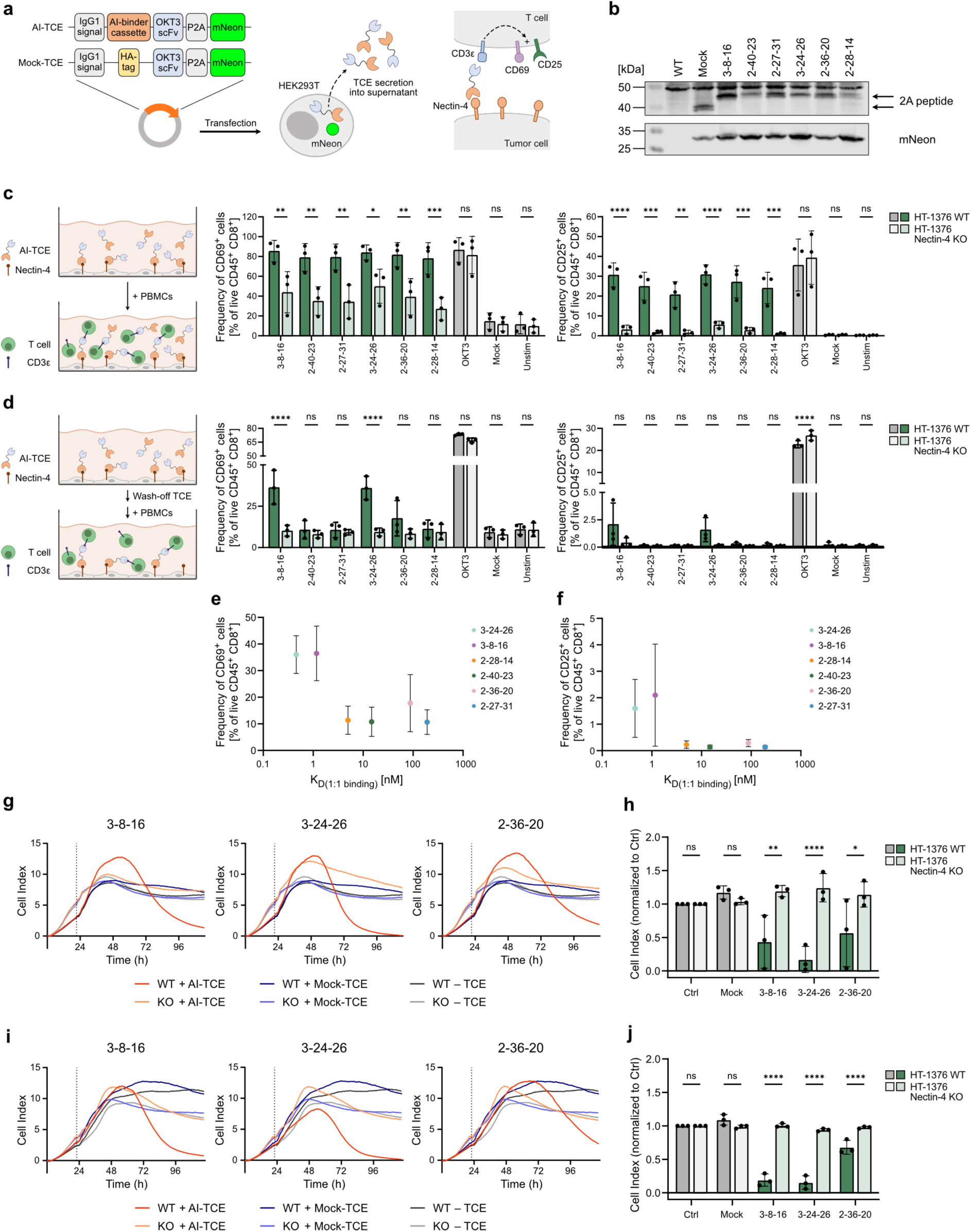
Functional assessment of Nectin-4 AI-minibinders in a translational application as bispecific T cell engager. **a)** Scheme illustrating the T cell engager (TCE) construct. AI-TCE constructs were transiently transfected into HEK293T cells and secreted into the supernatant. **b)** Western blot analysis showing the expression of TCE constructs in HEK293T cell lysates after transient transfection. **c)** Cartoon (left) showing the experimental set-up. Quantification of CD69^+^ CD8^+^ cells (middle) and CD25^+^ CD8^+^ cells (right) after 48 h co-culture of HT-1376 WT and Nectin-4 KO cells with human PBMCs and AI-TCEs or mock-TCE. Stimulation with an OKT3 antibody was used as positive control. **d)** Cartoon (left) showing the experimental set-up. Quantification of CD69^+^ CD8^+^ cells (middle) and CD25^+^ CD8^+^ cells (right) after 48 h co-culture of HT-1376 WT and Nectin-4 KO cells with human PBMCs and AI-TCEs or mock-TCE. Target cells were pre-incubated with TCEs, followed by the removal of the supernatant and an additional washing step. Stimulation with an OKT3 antibody was used as positive control. **e-f)** Dot plot showing the correlation between frequency of CD69^+^ CD8^+^ cells and CD25^+^ CD8^+^ cells from experiment described in d and their binding affinities (K_D_) determined by SPR. **g)** Representative curves showing co-cultures of human PBMCs with HT-1376 WT or Nectin-4 KO cells and AI-TCEs or mock-TCE measured as cell index value. Dashed line indicates the timepoint of effector addition. **h)** Quantification of the cell index value of experiment described in g at timepoint 72 h after effector addition. Cell index value was normalized to the control condition (co-culture without TCE). **i)** Representative curves showing co-cultures of human PBMCs with HT-1376 WT or Nectin-4 KO cells and AI-TCEs or mock-TCE measured as cell index value. TCE supernatant was pre-incubated on target cells and removed before effector cells were added. Dashed line indicates the timepoint of effector addition. **j)** Quantification of the cell index value of experiment described in i at timepoint 72 h after effector addition. Cell index value was normalized to the control condition (co-culture without TCE). Data are shown as mean ± SD performed in biological triplicates (n = 3). ns – non-significant; *p<0.05; **p<0.01; ***p<0.001; ****p<0.0001; two-way ANOVA with Šídák’s multiple comparisons test.

To generate Nectin-4-specific TCEs, we inserted several different Nectin-4 minibinder cassettes into the corresponding binding arm position of the TCE backbone. Secretion of the Nectin-4 TCE into the culture supernatant after plasmid transfection of HEK293T cells was confirmed by Western blot (**Figure 5b**). As a control, we generated a non-targeting mock-TCE by introducing a HA-tag in place of the Nectin-4 minibinder at the corresponding binding arm position of the TCE backbone. For the functional assays, we implemented two protocol variants. In variant 1, human peripheral blood mononuclear cells (PBMCs) were co-cultured with HT-1376 WT or Nectin-4 KO cells in the presence of TCE-containing HEK293T supernatant, that the TCE remained present throughout the assay (**Figure 5c**). HT-1376 Nectin-4 KO cells were included as matched specificity controls. Confirming successful target and T cell engagement, all Nectin-4 minibinder TCEs activated both CD8^+^ and CD4^+^ T cells in a target-specific manner, as indicated by surface upregulation of CD69 and CD25, two widely used T cell activation markers (**Figure 5c**, **Supplementary Figure 4a**)^10,20^. The fraction of CD69^+^ and CD25^+^ T cells was significantly higher in co-cultures with HT-1376 WT cells than with HT-1376 Nectin-4 KO cells. The bivalent OKT3 monoclonal antibody served as a positive control, highlighting that multivalent clustering of CD3 on T cells promotes activation, whereas the mock-TCE containing a monovalent OKT3 scFv served as a negative control.

Because baseline T cell activation on HT-1376 Nectin-4 KO cells was higher than in mock-TCE or no-TCE controls (**Figure 5c**), we tested a second protocol variant. In this setup, HT-1376 WT or HT-1376 Nectin-4 KO cells were first incubated with TCE-containing supernatant and then washed to remove unbound TCEs. PBMCs were subsequently added to the target cells, and activation was assessed as described above (**Figure 5d, Supplementary Figure 4b**). We reasoned that this approach provides a more stringent test of target-dependent T cell engagement, as only TCEs pre-bound to the cancer cell surface remains available to recruit and activate T cells.

Indeed, this pre-incubation and wash protocol completely abolished background T cell activation. Under these more stringent conditions, only two Nectin-4 minibinder TCEs (3-8-16 and 3-24-26) elicited statistically significant, target-specific T-cell activation, as indicated by CD69 and CD25 upregulation (**Figure 5d**). The 2-36-20 Nectin-4 TCE showed a similar trend but did not reach statistical significance. Notably, 3-8-16 and 3-24-26 corresponded to the highest-affinity minibinders identified above (K_D_ = 1.2 nM and 0.5 nM, respectively), and 3-8-16 also performed best in the quattrobinder approach (**Figure 5e, f**). Together, these results indicate that high-affinity target binding by AI-minibinders is critical for optimal TCE activity. Finally, we selected the top three Nectin-4 minibinder TCE constructs for dynamic target cell killing assays using the xCELLigence platform, a label-free, real-time cell analysis system that monitors living cells by measuring electrical impedance across microelectrodes. Again, we performed these experiments using both assay variants, co-culture with continuous TCE exposure (**Figure 5g, h, Supplementary Figure 4c**) and a pre-incubation and wash setup in which target cells were incubated with TCEs, washed, and then combined with PBMCs (**Figure 5i, j, Supplementary Figure 4d**). Consistent with the T cell activation assays, the 3-8-16 and 3-24-26 Nectin-4 TCEs mediated efficient, target-specific killing of HT-1376 WT cells, while sparing HT-1376 Nectin-4 KO cells. The 2-36-20 Nectin-4 TCE also induced specific killing, albeit to a lesser extent, in line with its lower binding affinity. In summary, our data show that Nectin-4 minibinders can be converted into potent, specific Nectin-4-targeting biologics, as demonstrated here with bispecific TCE constructs.

## Discussion

The AI-revolution has catapulted structural biology into an era where the reliable prediction of protein structures from their amino acid sequence^1,2,8^ and the design of completely artificial proteins are almost commonplace^4–7^. Multiple success stories have shown that it is now possible to design small proteins that bind to a target protein^4,5^, albeit with substantially varying success rates^10^. Here we set out to create AI-minibinders against the cancer target Nectin-4^13,14^, which in our hands turned out to be a particularly challenging target for the RFdiffusion algorithm. This is most likely due to the biophysical properties of its molecular surface. Nectin-4 lacks a pronounced hydrophobic patch, a feature that boosts design success in PD-L1, a structurally related protein that has become a benchmark target for protein design algorithms^4,5^.

Here, we show that combining a genetic algorithm with protein design algorithms can markedly improve AI-minibinder designs and increase experimental success rates for difficult targets. In fact, functional Nectin-4 AI-minibinders were obtained only after GA optimization, yielding highly specific binders with low nanomolar, down to subnanomolar affinities. Notably, none of the initial seed designs were recovered as functional binders in our mammalian cell-surface display screen. While we cannot exclude that individual seed designs might show measurable binding when tested one by one, the overall results clearly indicate that GA optimization substantially enhanced binder performance and the efficiency of their discovery. We also cannot exclude that other protein binder design pipelines, such as BindCraft^5^ or BoltzGen^7^, which were not tested here, may generate Nectin-4 minibinders more efficiently than RFdiffusion. Regardless, our GA optimization strategy is agnostic to the upstream design method and can readily use their outputs as seed designs in a highly compute-efficient manner. Notably, our results show that the performance of AI-minibinders with regard to a specific application can vary drastically. Our GA can very efficiently diversify a small set of seed designs, increasing the chances to identify an AI-minibinder that is suitable for a specific application.

Our work has focused on experimentally validating Nectin-4 minibinders generated by GA optimization using RFdiffusion’s partial diffusion feature, demonstrating that this strategy can serve as a powerful diversification component of evolutionary algorithms. In comparison, sequence-based optimization methods such as option II in Figure 1c offer complementary strengths^6^, most notably fine-grained control over the mutational space beyond point substitutions, including insertions, deletions, and crossover-like recombination events, while offering the possibility to easily mask certain regions of the design scaffold and add custom penalties, for instance to enforce an epitope of interest. However, for compact 3-helical AI-minibinders, these operations can be a double-edged sword, as indels and especially recombination events are more likely to disrupt the fold compared to simple point mutations. Thus, expanding the diversity of explored variants comes with a trade-off between broader sequence exploration and an increased risk of structure-destabilizing changes, and these parameters need to be carefully tested for each design problem.

We think that due to its similarity to natural evolution, the technique is very approachable for non-specialists. It is easy to adapt the parameters without the need to have a deep understanding of neural network architecture. While we have applied the technology to the optimization of protein–protein interactions, it can in principle be adapted to the redesign of protein–small molecule complexes, since algorithms such as Chai-1^6^, AF3^2^, and RoseTTAFold^3^ can be applied to such problems.

Target specificity is the decisive criterion for AI-minibinders, as high affinity is of limited value without unambiguous target engagement in a cellular context. To enable stringent specificity assessment, we employed the quattrobinder format for flow cytometry, which provides a robust readout of recognition of cell-surface-presented target protein^10^. Using isogenic wild-type and *NECTIN4*-knockout cells, we showed that the minibinders selectively stain Nectin-4-expressing cells, with knockout signals at background level, thus confirming target-specific binding. This emphasis on specificity is particularly important for *de novo*-designed proteins, which lack immunological selection and may therefore carry a higher risk of off-target interactions. However, cross-reactivity is also well documented for conventional immunization-derived antibodies, underscoring that specificity remains a key concern across binding modalities^24,25^.

Nectin-4 is the target of the approved antibody-drug conjugate (ADC) enfortumab vedotin (EV) and is highly expressed across several solid tumor types, particularly urothelial carcinoma of the bladder^15,13^. Clinical benefit from EV in patients with advanced metastatic UC is associated with high membranous Nectin-4 expression and amplification of the *NECTIN4* locus^13,14^. Beyond Nectin-4-directed ADCs, such as EV, multiple Nectin-4-targeting TCEs are in development and span diverse scaffolds, including bispecific antibodies^26^, multivalent VHHs (variable domains of heavy chain-only antibodies), and cyclic peptide formats^27^. Here, we demonstrate that specific, high-affinity Nectin-4 TCEs can be generated rapidly using *de novo* protein design, illustrating how these technologies may accelerate the development of next-generation cancer biologics. AI-minibinders offer several potential advantages, including their small size, which may enable improved tissue penetration, as well as facile engineering into multispecific formats and optimization of biophysical properties. To advance these candidates towards clinical application, however, comprehensive evaluation of key translational parameters, including safety and toxicity profiles, immunogenicity risk, pharmacokinetics, and biodistribution, is essential to establish their therapeutic viability and support progression into clinical testing.

In summary, advances in AI-enabled structural biology have made *de novo* binder generation increasingly accessible, yet performance remains strongly dependent on the target and on the specific design pipeline. Here, we introduce a versatile optimization algorithm that couples AI-derived seed designs to iterative, evolutionary refinement, enabling efficient exploration of sequence space through complementary diversification strategies, while using interaction-based scoring to prioritize improved variants. This framework consistently enhances candidate quality, developability and can be readily extended to optimize additional biophysical properties, providing a broadly applicable advance for accelerating therapeutic protein engineering.

## Online Methods

### Nectin-4 minibinder pool composition

Nectin-4 minibinder seed designs and GA-based refinement with option I (partial diffusion feature of RFdiffusion) were computed on local servers hosted by the Institute for Experimental Oncology (IEO) and the Institute of Structural Biology (ISB) at the University Hospital Bonn, Medical Faculty of the University of Bonn. The server used for the design tasks (AS -4125GS-TNRT, 2x AMD Genoa 9334 32C CPU, 768GB RAM) is equipped with NVIDIA RTX A5000 GPUs. In total, we performed 5 GA refinement runs. Run 1 (input: 12 Nectin-4 minibinder seed designs from 20,000 designs without predefined hotspots on Nectin-4); run 2 (input: 104 Nectin-4 minibinder seed designs from 20,000 designs with predefined hotspots on Nectin-4); run 3-5 top candidates from run 1 and 2. From all 5 GA runs, we collected the top-scoring and unique minibinder based on pAE interaction scores to obtain a GA-refined pool of in total 1876 minibinders. We also included the 12 and 104 seed designs resulting in a final pool size of 1992 minibinders that fits the tier 4 size (1,001-2,000) for DNA oligo pools from Twist Biosciences. PDB files, amino acid sequences, reverse translated DNA sequences and metrics (pAE scores) will be available via Zenodo repository (DOI: https://doi.org/10.5281/zenodo.18732269).

### GA-based refinement with Chai-1

GA-based minibinder refinement with option II (Chai-1) was computed on the Marvin HPC at the University of Bonn, Germany. Per GA run, we used one node of the scalable GPU applications (SGPU) partition with the following specifications: CPU: 2 × AMD EPYC “Milan” 64-core/128-thread 2.00GHz; RAM: 1024GB DDR4 3200MHz; GPU: 4 × NVIDIA A100 80GB (connected via NVLink within the node). For each refinement group, we performed 10 independent refinement runs with 50 generations and a pool size of 500. The point mutation rate was set to 5%; the indel and cross-over rates were set to 0%. For GA-refinement runs with binder pTM penalty, the pTM threshold was set to 0.9 and the weights were increased from 1 to 3. CIF files, amino acid sequences and metrics will be available via Zenodo repository (DOI: https://doi.org/10.5281/zenodo.18735047).

### DNA sequences

Primer sequences are provided in Supplementary Table 1.

### Library cloning

The obtained oligo pool from Twist Biosciences was amplified with PCR (primers: primers pool-amplification_fwd, primers pool-amplification_rev) for 20 cycles. A lentiviral backbone allowing the presentation of the minibinders in a chimeric antigen receptor-like construct was used and has been already previously described^10^. Backbone and amplified oligo pool were both pre-digested with Esp3I (Thermo Scientific, #FD0454) and cloned by a Golden Gate assembly approach. After ligation, the library was transformed into Endura electrocompetent *E. coli* (LGC Biosearch Technologies, #60242-1) by electroporation in 0.1 cm electroporation cuvettes (Bio-Rad, #1652089) with the following pulse conditions: 1800 V, 10 µF, 600 Ω. All bacteria from one reaction were resuspended in recovery medium (LGC Biosearch Technologies, #F98226-1) and cultured in a shaking incubator for 1 h at 37 °C and 300 rpm. The pre-culture was then transferred into 450 mL LB medium supplemented with 100 µg/mL ampicillin and incubated overnight at 37 °C with shaking at 180 rpm. The DNA was isolated using the PureLink™ HiPure Plasmid Midiprep Kit (Invitrogen, #K210005) according to manufacturer’s protocol. Coverage and representation of the AI-minibinder library was determined by next-generation sequencing (primers: NGS_cell-display_fwd, NGS_cell-display_rev, followed by a second PCR with barcode primers) using the Illumina MiSeq system.

### Cell culture

The human cell lines HEK293T (embryonic kidney, CVCL_0063), HT-1376 (bladder carcinoma, CVCL_1292), and HT-1376 Nectin-4 KO were cultivated in DMEM-GlutaMAX^TM^ (Gibco, #61965059) supplemented with 10% (v/v) of heat-inactivated FCS (Gibco, #A5256801) and 100 U/mL Penicillin-Streptomycin (Gibco, #15140163). The human cell lines RT4 (bladder carcinoma, CVCL_0036) and T24 (bladder carcinoma, CVCL_0554) as well as the Chinese hamster cell lines CHO-K1 (ovary, CVCL_0214), and CHO-Nectin-4 were cultivated in RPMI-GlutaMAX^TM^ (Gibco, #61870044) supplemented with 10% (v/v) of heat-inactivated FCS (Gibco, #A5256801) and 100 U/mL Penicillin-Streptomycin (Gibco, #15140163). Primary human cells were cultivated in RPMI-GlutaMAX^TM^ (Gibco, #61870044) supplemented with 10% (v/v) of heat-inactivated FCS (Gibco, #A5256801), 100 U/mL Penicillin-Streptomycin (Gibco, #15140163), 1 mM sodium pyruvate (Gibco, #11360088), 1× MEM non-essential amino acids (Gibco, #11140050), 10 mM HEPES (Carl Roth, #9157.1), 55 µM β-mercaptoethanol (Gibco, #21985023). All cell lines were cultured at 37 °C in 5% CO_2_ and regularly tested for mycoplasma contamination by PCR.

### Lentiviral transduction of AI-minibinder library

For lentiviral particle production, 1 × 10^6^ HEK293T cells were transiently transfected with 2 µg DNA of the AI-minibinder library plasmid, 0.65 µg DNA of lentiviral packaging plasmid psPAX2 (Addgene, #12260) and 0.35 µg DNA of lentiviral envelope plasmid pMD2.G (Addgene, #12259) in HEPES-buffered saline (HBS) and CaCl_2_. The medium was replaced twice, after 16 h and after 24 h, respectively. For the infection, the supernatant containing viral particles was collected 40 h post-transfection, filtered through 0.45 µm syringe filters and diluted 1:10 in normal medium. Previously plated 2 × 10^6^ HEK293T cells were infected with the diluted viral supernatant. After 48 h, transduced cells were selected by using medium supplemented with 1 µg/mL of puromycin.

### Screening using mammalian cell-surface display of library and cell sorting

HEK293T cells containing AI-minibinder library were harvested and stained according to standard flow cytometry protocols. Per sample, 4 × 10^6^ cells were blocked with human Fc receptor blocking solution (Biolegend, #422302) diluted 1:200 in PBS and afterwards incubated with 10 µg/mL of recombinant, Fc-tagged Nectin-4 protein from human, murine, rat, and cynomolgus origin (ACROBiosystems, Supplementary Table 1) in FACS buffer (PBS, 2% (v/v) FCS, 2 mM EDTA). Secondary staining was performed with 0.25 µg/mL of APC-labelled anti-human IgG Fc antibody (Biolegend, #410712, clone M1310G05). Cells were processed through a Sony MA900™ fluorescence-activated cell sorter and the APC positive fraction was sorted. The collected cells were lysed with direct lysis buffer (1 mM CaCl_2_, 3 mM MgCl_2_, 1 mM EDTA, 1% (v/v) Triton X-100, 10 mM Tris pH 7.5) containing 0.2 mg/mL proteinase K and incubated for 15 min at 56 °C and for another 15 min at 95 °C afterwards. Samples were prepared for next-generation sequencing by PCR amplification (primers: NGS_cell-display_fwd, NGS_cell-display_rev) and barcoded in a second PCR amplification step (primers: barcode primers) using the NEBNext High-Fidelity 2× PCR MasterMix (NEB, #M0541). Subsequently, all samples were sequenced with the Illumina MiSeq system. Reads were filtered for a Q-score>30, matched to a reference containing all AI-minibinder sequences from the library and enrichments of AI-minibinders were analyzed (https://jsb-lab.bio/ai-binders/).

### Generation of monoclonal cell lines and overexpression cell lines

A CRISPR-Cas9-mediated knockout of *NECTIN4* was previously generated in HT-1376 cells prior to this study^14^. Cells of the polyclonal HT-1376 Nectin-4 KO cell line were stained 1:100 with APC-labelled anti-human Nectin-4 antibody (Miltenyi Biotec, #130-116-028, clone REA967). Further selection of the negative population was performed by fluorescence-activated cell sorting with a BD FACSAria™ III Cell Sorter. Absence of Nectin-4 expression was validated with flow cytometry afterwards and indel frequency of *NECTIN4* was analyzed with targeted amplicon next-generation sequencing. Therefore, genomic DNA from the sorted cell line was isolated using the Nucleo Spin Tissue Kit (Macherey & Nagel, #740952.250). The gene region of interest was amplified with a first PCR (primers: MiSeq_hsNECTIN-4-KO1_fw, MiSeq_hsNECTIN-4-KO1_rev) and barcoded in a second PCR step (primers: barcode primers). All samples were sequenced with the Illumina MiSeq system and indel frequency was determined using a web-based tool (http://www.outknocker.org/outknocker2.htm)^26^. Experiments were performed with the sorted Nectin-4 KO cell line, if not otherwise stated. Polyclonal HT-1376 Nectin-4 KO cell line was harvested and plated into 96-well plates in a concentration of 0.7 cells/well. After formation of colonies from single cells, monoclonal cell lines were expanded and genomic DNA was isolated as aforementioned. Phenotype was screened by flow cytometry staining of human Nectin-4 (Miltenyi Biotec, #130-116-028, clone REA967) with a BD FACS Canto™ II analyzer and genotyping was performed as described above for the polyclonal culture.

To generate the human Nectin-4 overexpressing cell line CHO-Nectin-4, the coding sequence of *NECTIN4* was retrieved from cDNA of MCF-7 cells (breast cancer, CVCL_0031) by PCR (primers: hNectin4_HindIII_fwd, hNectin4_SalI_rev) and cloned into pRP233 plasmid (derived from Addgene plasmid, #41841). HEK293T cells were transiently transfected with HEPES-buffered saline (HBS) and CaCl_2_, 2 µg of the generated plasmid, 2 µg of the retroviral packaging plasmids gag-pol and 220 ng of pCMV VSV-G. The medium was changed 16 h after transfection and the virus containing supernatant was collected 40 h post-transfection and filtered with 0.45 µm syringe filters. CHO-K1 cells were incubated with the viral particles and successfully transduced cells were selected 48 h after infection with medium containing 25 µg/mL puromycin. Further selection was performed by fluorescence-activated cell sorting with a BD FACSAria™ III Cell Sorter. The cells were stained 1:100 with APC-labelled anti-human Nectin-4 antibody (Miltenyi Biotec, #130-116-028, clone REA967) and the positive population was sorted.

### Cloning and validation of individual AI-minibinder hits

Individual AI-minibinder sequences were ordered as gene fragments (Twist Biosciences) and cloned into the lentiviral backbone for the mammalian cell-surface display using a Golden Gate assembly with Esp3I (Thermo Scientific, #FD0454). Successful cloning was confirmed via sanger sequencing (primer: Sanger_minibinder_fwd). 300,000 HEK293T cells were transiently transfected with 1 µg of the AI-minibinder constructs using HEPES-buffered saline (HBS) and CaCl_2_. Medium was changed 16 h after transfection. After 40 h post-transfection, cells were collected and stained with a standard flow cytometry protocol. Firstly, cells were blocked with human Fc receptor blocking solution (Biolegend, #422302) diluted 1:200 in PBS and incubated with Zombie Violet™ fixable dye (Biolegend, #423114) diluted 1:2,500 in PBS. Then, samples were stained with 10 µg/mL of either human, murine, rat, or cynomolgus recombinant Fc-tagged Nectin-4 proteins (ACROBiosystems, Supplementary Table 1). A secondary staining was performed with 0.25 µg/mL APC-labelled anti-human IgG Fc antibody (Biolegend, #410712, clone M1310G05). To control for surface expression, samples were stained with 2 µg/mL APC-labelled anti-HA.11 epitope tag antibody (Biolegend, #901523, clone 16B12). Samples were acquired at the BD FACS Canto™ II analyzer and downstream analysis was performed with FlowJo. For each sample, the APC geometric mean fluorescence intensity of protein binding was normalized to the respective HA.11 epitope tag signal.

### Cloning of AI-minibinders for bacterial expression

AI-minibinder coding sequences were amplified from the lentiviral backbone of the mammalian cell-surface display system and subcloned into the pEHISTEV backbone for expression in *E. coli*^17^. Cysteine point mutations were introduced by primer-directed mutagenesis following the method by Liu and Naismith^28^.

### Expression and purification of AI-minibinders

Recombinant 6xHis-tagged AI-minibinders were expressed and purified as described previously^9^. In brief, *E. coli* BL21 (DE3) bacterial cells were transformed with the respective constructs and grown to saturation at 37 °C overnight. The saturated culture was used for inoculation of main cultures, which were grown at 37 °C to an OD_600_ of 0.8. IPTG was added to a final concentration of 0.5 mM and proteins were expressed at 18 °C for 16 h. Cells were harvested at 4,000 × g for 25 min and flash-frozen in liquid nitrogen or subjected to immediate cell lysis.

For affinity chromatography, cell pellets were resuspended in lysis buffer (20 mM Tris pH 8.0, 0.5 M NaCl, 20 mM imidazole) and lysed by sonication. The lysate was cleared by centrifugation at 25,000 × g for 45 min at 10 °C. The supernatant was filtered through a 0.8 µm pore size membrane and incubated with Ni^2+^-NTA resin for 2 h at 4 °C or room temperature. The resin was washed extensively with lysis buffer and proteins were eluted by addition of lysis buffer supplemented with 0.5 M imidazole. For cysteine mutants, the lysis buffer additionally contained 5 mM β-mercaptoethanol.

Following affinity chromatography, proteins were further purified by size exclusion chromatography (SEC) using a HiLoad 16/600 Superdex 75 pg column (Cytiva) equilibrated in SEC buffer (20 mM Tris pH 8.0, 250 mM NaCl). For cysteine mutants, the SEC buffer additionally contained 2 mM TCEP.

### Biotinylation of AI-minibinders

AI-minibinders containing cysteine point mutations were biotinylated using the cysteine-reactive maleimide-PEG_2_-biotin label (EZ-Link, ThermoFisher). Before biotinylation, AI-minibinders were buffered into 20 mM Tris pH 8.0, 250 mM NaCl using 7 K MWCO Zeba Spin columns to remove TCEP from the previous SEC purification step. The label was added at a 10-fold molar excess and incubated overnight at 4 °C. Excess label was removed by desalting using 7 K MWCO Zeba Spin columns.

### Quattrobinder assembly and validation by flow cytometry

In *E. coli* expressed and biotinylated AI-minibinder monomers were coupled to Alexa Fluor™ 647-conjugated streptavidin (Invitrogen, #S21374) to form tetravalent quattrobinders. Therefore, streptavidin was step-wise added to the minibinders until a total molar ratio of 4:1 was reached. Every step was followed by 10 min incubation at room temperature. Assembled quattrobinders were validated by flow cytometry. A concentration titration experiment with 150 nM, 75 nM and 37 nM of quattrobinder solution was conducted to identify the optimal concentration. Target and non-target expressing cell lines were harvested and blocked with human Fc receptor blocking solution (Biolegend, #422302) diluted 1:200 in PBS and incubated with Zombie Violet™ fixable dye (Biolegend, #423114) diluted 1:2,500 in PBS. The samples were stained with 75 nM of quattrobinder solution or as control with a conventional biotinylated anti-Nectin-4 antibody (Miltenyi Biotec, #130-116-100, clone REA967, 1:100 dilution) and a secondary staining step with 75 nM of Alexa Fluor™ 647-conjugated streptavidin (Invitrogen, #S21374) in FACS buffer (PBS, 2% (v/v) FCS, 2 mM EDTA). The samples were acquired with a BD FACS Canto™ II analyzer. Analyses of the experiments were performed with FlowJo.

### Cloning of expression plasmids for AI-minibinder TCE and their production

AI-minibinder sequences were ordered as individual gene fragments (Twist Biosciences) and cloned into the bispecific T cell engager (TCE) expression plasmid (Twist Biosciences) with a Golden Gate assembly using BbsI. The sequences of the TCE cassette were obtained from Sheperd et al.^18^ and modified as needed. Successfully cloned constructs were validated through Sanger sequencing (primer: Sanger_TCE_fwd). For the production of AI-minibinder TCEs into the supernatant, 1 × 10^6^ HEK293T cells were transiently transfected with HEPES-buffered saline (HBS) and CaCl_2_ and 2 µg of the respective AI-minibinder TCE construct. 16 h post-transfection, the medium was exchanged to RPMI-GlutaMAX^TM^ (Gibco, #61870044) supplemented with 10% (v/v) of heat-inactivated FCS (Gibco, #A5256801), 100 U/mL Penicillin-Streptomycin (Gibco, #15140163), 1 mM sodium pyruvate (Gibco, #11360088), 1× MEM non-essential amino acids (Gibco, #11140050), 10 mM HEPES (Carl Roth, #9157.1), 55 µM β-mercaptoethanol (Gibco, #21985023). HEK293T cell supernatant was collected 40 h after transfection and filtered through 0.45 µm syringe filters. The supernatant was either directly used in following experiments or stored at −20 °C.

### AI-minibinder TCE in activation co-culture assay with human PBMCs

Peripheral blood from healthy donors was provided by the Institute for Experimental Hematology and Transfusion Medicine at the University Hospital Bonn, Bonn, Germany. All study procedures were performed in compliance with relevant laws and institutional guidelines and have been approved by the competent Ethics Committee of the Medical Faculty of the University of Bonn. Written informed consent was received prior to participation from the blood donors. Peripheral blood mononuclear cells (PBMCs) were thawed and rested for 1 h at 37 °C with 10 µg/mL DNAse I (Roche, #10104159001). After resting, 1 × 10^6^ PBMCs were resuspended in 1 mL AI-TCE or mock-TCE supernatant with 100 U/mL rhIL-2 (PeproTech, #200-02) and co-cultured with previously plated HT-1376 WT or Nectin-4 KO target cells, 30,000 cells per 48-well, for 48 h at 37 °C. In a second experimental set-up, target cells were pre-incubated with filtered HEK293T supernatant containing AI-minibinder TCEs for 2 h at 37 °C. After incubation, supernatant was discarded and unbound TCEs were removed by washing with PBS before adding PBMCs to the target cells. Stimulation with 1 µg/mL of CD3 antibody (Biolegend, clone OKT3, #317325) was used as positive control. PBMCs after co-culturing were collected and blocked with human Fc receptor blocking solution (Biolegend, #422302) diluted 1:200 in PBS and incubated with Zombie Green™ fixable dye (Biolegend, #423112) diluted 1:1,000 in PBS. Flow cytometry staining of surface markers with all antibodies (CD45, Biolegend, #304032, clone HI30; CD3, BD Biosciences, #557832, clone SK7; CD8, Biolegend, #344756, clone SK1; CD4, BD Biosciences, #750975, clone OKT4; CD69, Biolegend, #310906, clone FN50; CD25, Biolegend, #302640, clone BC69) was performed at a 1:200 dilution in FACS buffer (PBS, 2% (v/v) FCS, 2 mM EDTA) and all samples were acquired with a 4-LaserCytek Aurora Spectral Flow Cytometer. FlowJo was utilized for downstream analyses.

### Western blots

HEK293T cells were transfected as described for the AI-minibinder TCE production and collected after harvesting the TCE supernatant. Urothelial carcinoma cells and transfected HEK293T cells were lysed using RIPA lysis buffer (150 mM NaCl, 50 mM TRIS-HCl pH 8.0, 1% (v/v) Nonidet P-40, 0.5% (w/v) Sodium deoxycholate, 0.1% (w/v) SDS) supplemented with protease and phosphatase inhibitors diluted 1:100 (Cell Signaling Technology, #5872). The total protein concentration was determined using the Bradford method with the Protein Assay Dye Reagent (Bio-Rad, #5000006). For each sample, 20 μg of protein were prepared in 6× Laemmli buffer (375 mM Tris pH 6.8, 12% (w/v) SDS, 47% (v/v) glycerol, 9.3% (w/v) DTT, 0.06% (w/v) bromophenol blue) and loaded on 12% or 15% polyacrylamide gels. After separation via SDS-PAGE, proteins were transferred to nitrocellulose blotting membranes (Cytiva, #10600004) in a wet blotting Mini Trans-Blot Cell system (Bio-Rad). The membranes were blocked with 5% (w/v) bovine serum albumin (BSA) (Carl Roth, #8076) in TRIS-buffered saline with 0.05% (v/v) Tween 20 (TBS-T) for 1 h at room temperature followed by incubation with primary antibodies overnight at 4 °C. All primary antibodies were diluted in blocking solution (5% (w/v) BSA in TBS-T) with the following dilutions: anti-2A peptide (Novus Biologicals, clone 3H4, 1:1000, #NBP2-59627), anti-mNeon (Cell Signaling Technology, polyclonal, 1:1000, #53061), anti-Nectin-4 (R&D Systems, polyclonal, 1:500, #AF2659), and anti-β-actin (Santa Cruz Biotechnology, clone C4, 1:1000, #sc-47778). After overnight incubation with primary antibodies, the membranes were washed with TBS-T and incubated with secondary antibodies for 1 h at room temperature. All secondary antibodies were used at 1:15000 diluted in blocking solution: IRDye 800CW Goat anti-Mouse (LICORbio, #926-32210), IRDye 680RD Goat anti-Rabbit (LICORbio, #926-68071), IRDye 680RD Donkey anti-Mouse (LICORbio, #926-68072), and IRDye 800CW Donkey anti-Goat (LICORbio, #926-32214). After incubation with secondary antibodies, the membranes were washed with TBS-T and the protein bands were imaged with the Odyssey CLx system (LICORbio).

### Measuring cytotoxicity of AI-minibinder TCEs

For this assay, the xCELLigence RTCA SP system (Agilent Technologies) was used with a protocol adapted from Kanemaru et al.^27^ In brief, HT-1376 WT or Nectin-4 KO cells were seeded in a density of 10,000 cells in 100 µL media per well in E-Plate 96 PET (Agilent Technologies, #300600910). Growth behavior was monitored by measuring the electrical impedance in 15 min intervals, which is indicated as cell index value. Target cell growth into a monolayer was determined for approximately 24 h. Co-culture of target cells and human PBMCs was set-up as described for the activation assay. Per well, 0.2 × 10^6^ PBMCs were resuspended in 200 µL of AI-TCE or mock-TCE with 100 U/mL rhIL-2 (PeproTech, #200-02) and added to the target cells. Similar to the activation assay, in the second experimental set-up, target cells were pre-incubated with TCE-containing supernatant for 2 h at 37 °C. Excess supernatant was removed and unbound TCEs were washed off before the addition of PBMCs. Cell index value was measured every 15 min for the following 96 h. Conditions of PBMCs alone and target cells alone were included as background controls.

### Protein thermal stability analyses

The thermal stability of purified AI-minibinders was analyzed by nanoscale differential scanning fluorimetry (nanoDSF) on a Prometheus NT.48 (NanoTemper) device. AI-minibinders were adjusted to a concentration of 10 µM in 20 mM HEPES pH 7.4, 150 mM NaCl. Protein denaturation and renaturation events were monitored by changes in internal fluorescence at wavelengths of 330 and 350 nm. The changes were recorded from 20 °C to 90 °C (unfolding) followed by 90 °C to 20 °C (refolding) at a rate of 1.5 °C/min and evaluated using the PR.ThermControl software (NanoTemper).

### Analytical size exclusion chromatography

Analytical size exclusion chromatography of AI-minibinders was performed on a Superdex Increase 75 3.2/300 column (Cytiva) equilibrated in 20 mM HEPES pH 7.4, 150 mM NaCl and run at a flow-rate of 0.05 mL/min using the Infinity II HPLC System (Agilent). A commercial SEC standard (BioRad, #1511901) was run as a molecular weight reference.

### Surface plasmon resonance measurements

Surface plasmon resonance (SPR) measurements were performed on a Series S CM5 sensor chip (Cytiva) using a Biacore 8K instrument (GE Healthcare) operated at 25 °C. The chip was equilibrated in running buffer (20 mM HEPES pH 7.4, 150 mM NaCl, 1 mM TCEP, 0.05% (v/v) Tween-20). The sensor chip surface was activated using a 1:1 (v/v) mixture of 0.1 M *N*-hydroxysuccinimide (NHS) and 0.1 M 3-(*N*,*N*-dimethylamino)propyl-*N*-ethylcarbodimide (EDC). For immobilization, 1 µM human Nectin-4 (1-336) in 10 mM sodium acetate buffer (pH 4.5) was injected over the surface of the second flow cell for 600 seconds at a flow-rate of 10 µL/min. Free binding sites on the surface were blocked with 1 M ethanolamine (pH 8.0) for 420 seconds at a flow-rate of 10 µL/min. Single-cycle kinetics were measured with injections of increasing concentration of the analyte of both flow cells. The association and dissociation times were set to 120 seconds and 600 seconds, respectively, at a flow-rate of 30 µL/min. For 2-28-14, a five-fold concentration series ranging from 0.00768 nM to 24 nM was used. For 3-8-16 and 3-24-26, a five-fold concentration series ranging from 0.0384 nM to 120 nM was used. For 2-27-31, 2-36-20, 2-35-9, and 2-40-23, a five-fold concentration series ranging from 4.8 nM to 15,000 nM was used.

### Expression and purification of human Nectin-4

Recombinant human Nectin-4 was expressed from a pcDNA3.1 (+) plasmid containing a HRV 3C protease-cleavable C-terminal 8xHis-Twin-Strep II tag. Expi293F cells were grown in FreeStyle 293F medium and passaged to fresh medium on the day before transfection at a density of 2 × 10^6^ cells/mL. The next day, the appropriate volume of culture was centrifuged (5 min, room temperature, 100 × g) and the cell pellet was resuspended in 52.5 mL fresh FreeStyle medium to a final density of 20 × 10^6^ cells/mL. 3 × 17.5 mL cell suspension was transferred to 3 TubeSpin® bioreactor 600. DNA was added to a final concentration of 1.5 µg/million cells and mixed. Immediately, 3 µg/million cells of a 1 mg/mL PEI MAX stock solution (Polyethylenimine HCL MAX, MW 40000, Polysciences) was added. The cultures were again swirled and then transferred to an incubator shaker with 8% CO_2_ for 1 h at 37 °C with agitation at 180 rpm (Minitron incubator, INFORS HT). At the end of the transfection phase, 332.5 mL of pre-warmed FreeStyle 293 medium was added to each bioreactor to give a density of 1 × 10^6^ cells/mL. The cells were incubated for 96 h (37 °C, 8% CO_2_, 70% humidity, 180 rpm). The protein-containing medium was harvested 4 days post-transfection (3200 × g, 20 min, 4 °C). 105 mL of 10× Strep tag buffer (500 mM TRIS-HCl pH 8, 1.5 M NaCl) and 2 mL of BioLock (Biotin blocking solution, IBA) were added and gently stirred for 10 min. The medium was filtered (pore size of 0.45 µm) prior to purification.

The filtered protein-containing medium was loaded on a 1 mL Strep-Tactin cartridge (IBA, #2-1237-001) equilibrated in wash buffer (50 mM TRIS-HCl pH 8.0, 150 mM NaCl), washed extensively with wash buffer and eluted in elution buffer (50 mM TRIS-HCl pH 8.0, 150 mM NaCl, 5 mM desthiobiotin) using a sample pump. The elution fractions were pooled and the C-terminal tag was removed by incubation with HRV 3C protease overnight at 4 °C. The next day, 10× His tag buffer (500 mM TRIS-HCl pH 8.0, 1.5 M NaCl, 200 mM imidazole) was added to the protein sample to a final concentration of 50 mM TRIS-HCl pH 8.0, 150 mM NaCl, 20 mM imidazole. For removal of the tag, 50 µL Ni^2+^-NTA agarose beads (QIAGEN) were added and incubated for 1 h at 4 °C on a rolling incubator. The sample was poured into a low volume Poly-Prep Column (Bio-Rad) and the flow-through was collected. The flow-through containing digested Nectin-4 was concentrated using an Amicon Ultra 4 mL Filter with 10 K cut-off (Merck Millipore). The protein was further purified on a Superdex 200 Increase 10/300 GL column (GE) equilibrated with SEC running buffer (20 mM HEPES pH 7.5, 100 mM NaCl, 10% (v/v) glycerol). The protein purity was confirmed by Coomassie-stained SDS-PAGE. Fractions were pooled and concentrated using an Amicon Ultra 4 mL Filter with 10 K cut-off (Merck Millipore) and flash frozen.

## Code availability

Code for NGS analysis and genetic algorithms will be available via GitHub.

## Data availability

PDB files, amino acid sequences, reverse translated DNA sequences and metrics (pAE scores) for the Nectin-4 minibinder pool will be available via Zenodo repository (DOI: https://doi.org/10.5281/zenodo.18732269). CIF files, amino acid sequences and metrics for the minibinder pool of GA-based refinement with Chai-1 will be available via Zenodo repository (DOI: https://doi.org/10.5281/zenodo.18735047).

## Acknowledgements

We would like to thank the Flow Cytometry Core Facility of the Medical Faculty at the University of Bonn for providing support and instrumentation. We would also like to thank the NGS Core Facility of the Medical Faculty at the University of Bonn for providing support and instrumentation. We would like to thank Johannes Siewert for generating the CHO-Nectin-4 overexpression cell line. We would like Yexin Xie and Kim Remans from the EMBL PepCore (Heidelberg) for recombinant Nectin-4 protein production and purification.

The authors gratefully acknowledge the access to the Marvin cluster of the University of Bonn and the support provided by the High-Performance Computing & Analytics Lab of the University of Bonn.

This project was supported by a consortial grant (PrepAIred - Precision targeting of pancreatic cancer by generative AI-based protein design) from the Deutsche Krebshilfe (German Cancer Aid) – project grant ID 70117114 – to M.I.T, T.B, J.L.S.-B., M.H. and G.H. For this project, J.S.B. was also supported – in part – by the Hans und Ria Messer Stiftung and the Gates Foundation.

M.H. was also supported – in part - by the Deutsche Krebshilfe (German Cancer Aid) project grant 70114292 (Excellence Program for established scientists), project grant 70116843 by the Deutsche Krebshilfe and within SFB 1399 by the Deutsche Forschungsgemeinschaft (DFG) – Project ID 413326622. E.B. was supported by the Deutsche Krebshilfe (German Cancer Aid) with a MD doctorate scholarship - grant 70113307 (Mildred Scheel Nachwuchszentrum). This project was in part supported by a consortial grant (TENACITY) funded under the ERA-Net JTC2021 TRANSCAN-3 initiative – project ID 01KT2301 – to T.B. and M.H.

M.H. is a member of the CANTAR project, which receives funding from the Netzwerke 2021 program, an initiative of the Ministry of Culture and Science of the State of North Rhine-Westphalia (the sole responsibility for the content of this publication lies with the authors). M.G., M.I.T, N.K., T.B, J.L.S.-B., M.H and GH are members and supported by ImmunoSensation^3^ – the immune sensory system – supported by the Deutsche Forschungsgemeinschaft under Germany’s Excellence Strategy EXC2151 - project ID 390873048.

## Author contributions

Acquisition of data: E.T., S.C.F., B.B., J.M.M., B.A.McE., M.C.R.Y., D.F., L.W.

Data analysis and visualization: E.T., S.C.F., T.N.K., M.H., G.H.

Resources: M.G., J.O., H.R., M.I.T., N.K., T.B., A.H., J.L.S.-B., M.H., G.H., E.B.

Cartoons, icons: E.T.

Protein design: M.H., G.H.

Coding: M.H., G.H., T.N.K., B.B., J.L.S.-B.

Conceptualization: M.H., G.H.

Reviewing and editing: B.B., J.M.M., B.A.McE, M.C.R.Y., M.G., J.O., H.R., M.I.T., N.K., T.B., J.L.S.-B.

Writing: M.H., G.H, E.T., S.C.F.

All authors read and approved the final manuscript.

## Supplementary figures

**Supplementary Figure 1:**
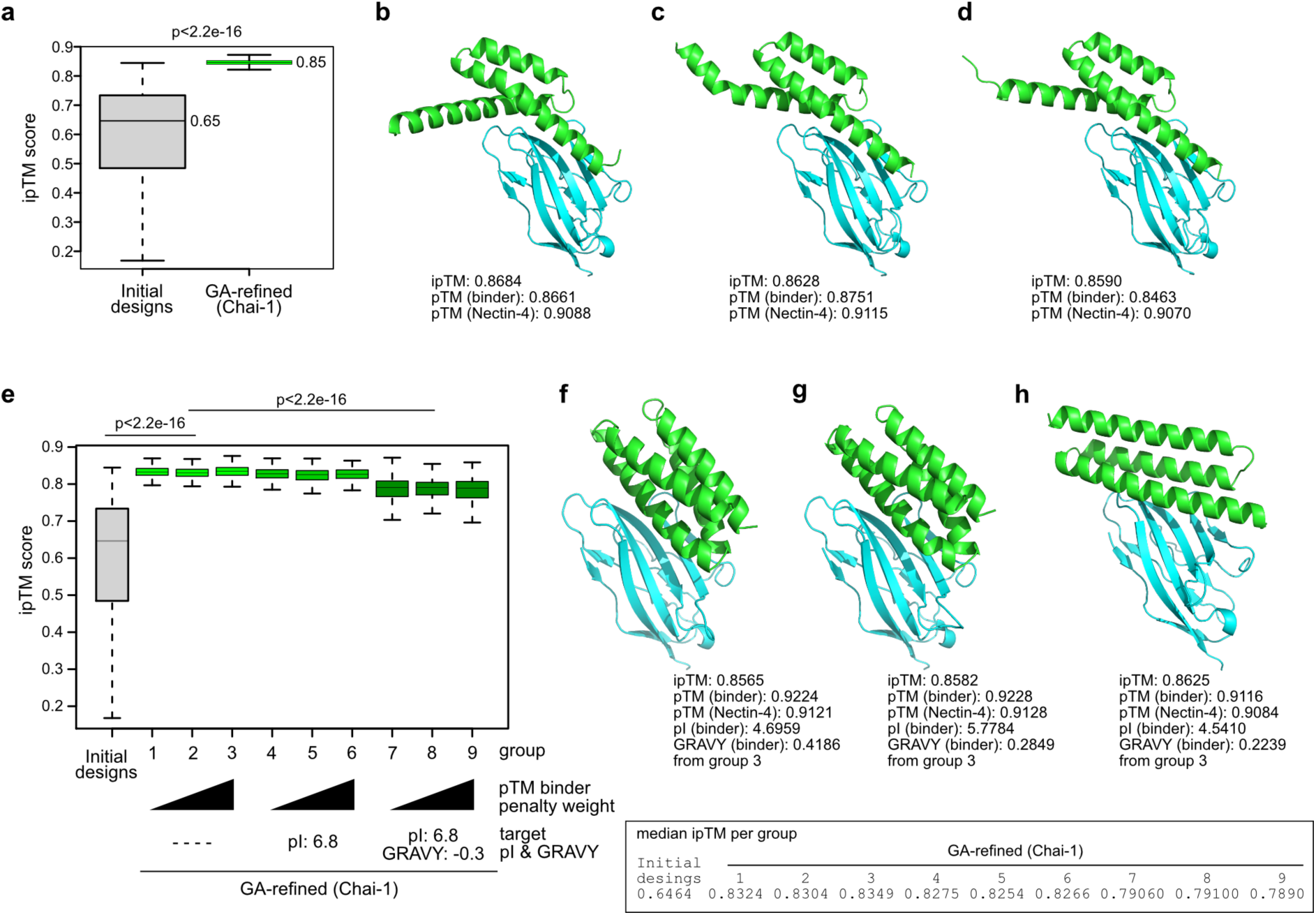
Refinement of Nectin-4 minibinders by option II genetic algorithm with Chai-1 enabled sequence editing. **a)** Boxplot showing differences in ipTM scores between Nectin-4 minibinder initial designs and option II GA-redefined designs. Median ipTM scores are indicated. Two-sided Wilcoxon test. **b-d)** Three examples of predicted structures of option II GA-refined Nectin-4 minibinders in complex with Nectin-4. Respective ipTM and pTM scores are indicated. **e)** Boxplot showing differences in ipTM scores between Nectin-4 minibinder initial designs and option II GA-redefined designs with implemented pTM binder penalty and with or without target isoelectric point (pI) and hydrophobicity (GRAVY score). Pairwise two-sided Wilcoxon test with Benjamini & Hochberg correction for multiple testing. **f-h)** Three examples (from group 3) of predicted structures of option II GA-refined Nectin-4 minibinders with pTM penalties. Respective ipTM and pTM scores are indicated.

**Supplementary Figure 2:**
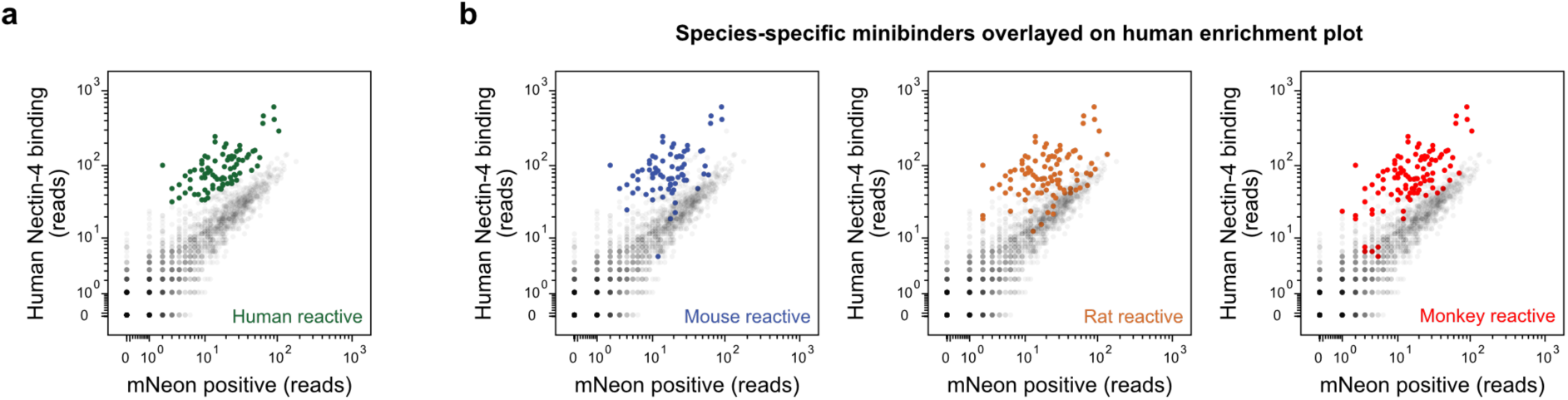
Identification of cross-species reactive AI-minibinders. **a)** Scatter plot showing the enriched minibinder population for human Nectin-4 (green dots). **b)** Scatter plots of the enriched minibinder population detected from the screen with murine, rat and monkey Nectin-4 Fc-fusion proteins overlaid into the human Nectin-4 enrichment plot (colored dots).

**Supplementary Figure 3:**
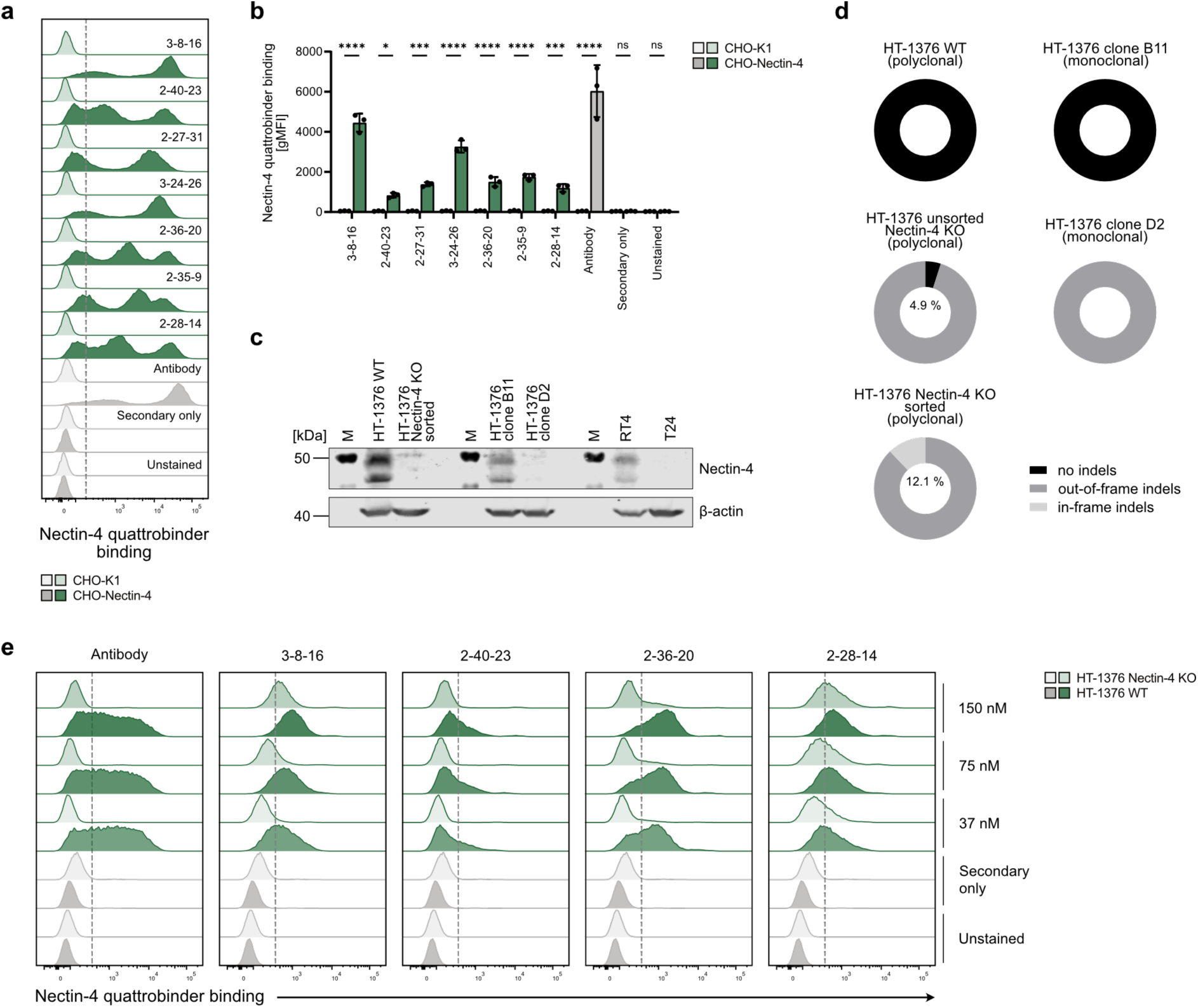
Optimization of quattrobinder staining and validation of urothelial cancer cell lines. **a)** Representative histograms showing flow cytometry analyses of AF647-conjugated Nectin-4 quattrobinders in comparison to a conventional biotinylated Nectin-4 antibody with a secondary AF647-streptavidin staining on target overexpressing CHO-Nectin-4 cells and parental CHO-K1 cells. **b)** Quantification of the experiment described in a. Data are shown as mean ± SD performed in biological triplicates (n = 3). **c)** Western blot analysis displaying Nectin-4 expression levels of various urothelial carcinoma cell lines used in this study. **d)** Next-generation sequencing analysis of genomic DNA from polyclonal HT-1376 WT and Nectin-4 KO cells prior and after cell sorting as well as from monoclonal HT-1376 cell lines. **e)** Histograms showing a concentration titration of the conventional biotinylated Nectin-4 antibody with secondary AF647-streptavidin staining and four AF647-conjugated Nectin-4 quattrobinders (3-8-16, 2-40-23, 2-36-30, 2-28-14) on HT-1376 WT and the corresponding knockout cell line (HT-1376 Nectin-4 KO). ns – non-significant; *p<0.05; ***p<0.001; ****p<0.0001; two-way ANOVA with Šídák’s multiple comparisons test.

**Supplementary Figure 4:**
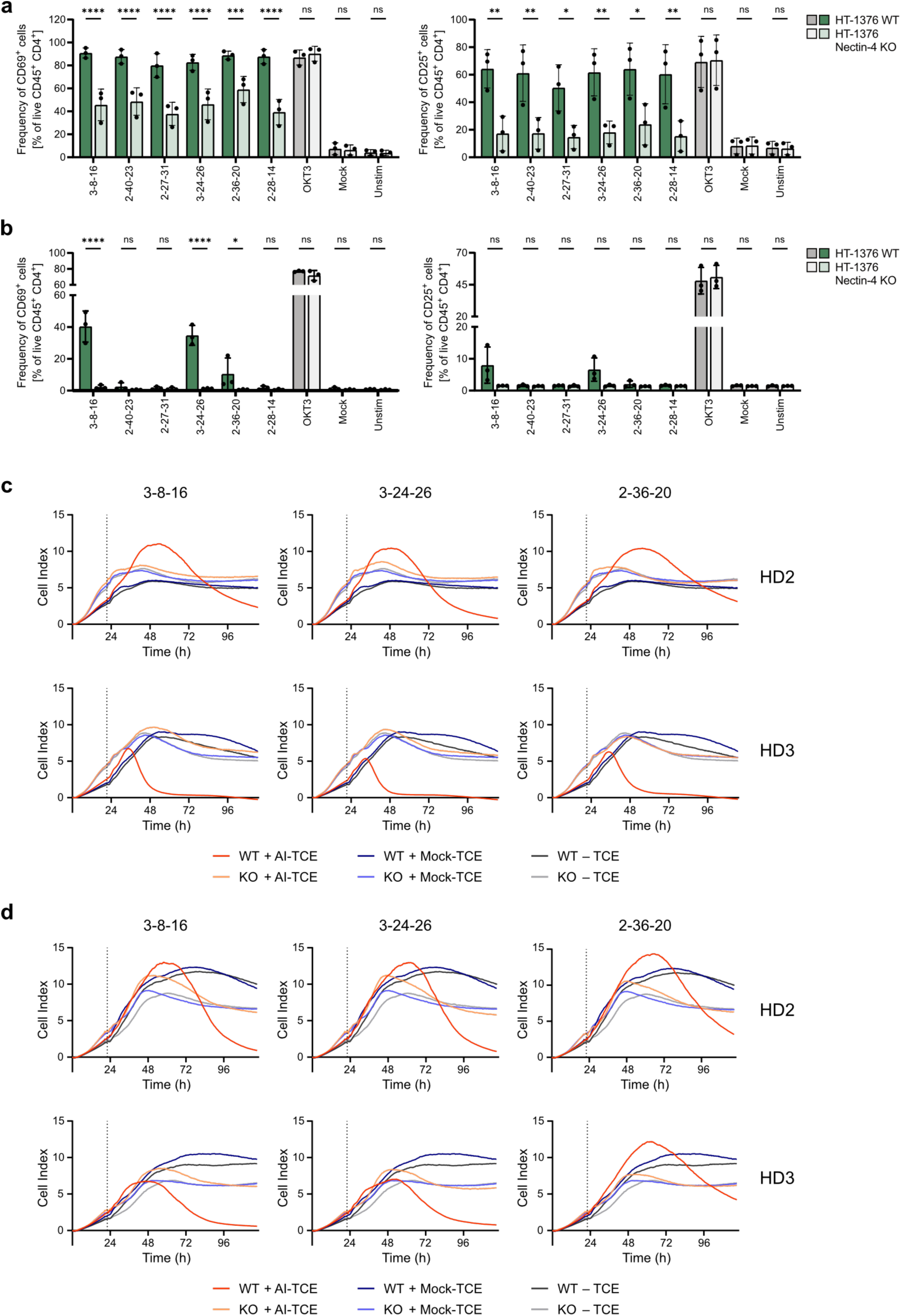
Successful application of Nectin-4 AI-minibinders as bispecific T cell engager. **a)** Quantification of CD69^+^ CD4^+^ cells (left) and CD25^+^ CD4^+^ cells (right) after 48 h co-culture of HT-1376 WT and Nectin-4 KO cells with human PBMCs and AI-TCEs or mock-TCE. Stimulation with an OKT3 antibody was used as positive control. **b)** Quantification of CD69^+^ CD4^+^ cells (left) and CD25^+^ CD4^+^ cells (right) after 48 h co-culture of HT-1376 WT and Nectin-4 KO cells with human PBMCs and AI-TCEs or mock-TCE. Target cells were pre-incubated with TCEs, followed by the removal of the supernatant and an additional washing step. Stimulation with an OKT3 antibody was used as positive control. **c)** Curves of replicate 2 and 3 showing co-cultures of human PBMCs with HT-1376 WT or Nectin-4 KO cells and AI-TCEs or mock-TCE measured as cell index value. Dashed line indicates the timepoint of effector addition. **d)** Curves of replicate 2 and 3 showing co-cultures of human PBMCs with HT-1376 WT or Nectin-4 KO cells and AI-TCEs or mock-TCE measured as cell index value. TCE supernatant was pre-incubated on target cells and removed before effector cells were added. Dashed line indicates the timepoint of effector addition. Data are shown as mean ± SD performed in biological triplicates (n = 3). ns – non-significant; *p<0.05; **p<0.01; ***p<0.001; ****p<0.0001; two-way ANOVA with Šídák’s multiple comparisons test.

